# E74 like ETS transcription factor 3 (ELF3) is a negative regulator of epithelial-mesenchymal transition in bladder carcinoma

**DOI:** 10.1101/496646

**Authors:** Kirti Gondkar, Krishna Patel, Shoba Krishnappa, Akkamahadevi Patil, Bipin Nair, Gopinath Meenakshi Sundaram, Tan Tuan Zea, Prashant Kumar

## Abstract

Transcription factors are known to be commonly deregulated in various cancers. The E74 like ETS transcription factor 3 (ELF3) expression is restricted to epithelial tissue. In the present study, we evaluated the role of ELF3 in the pathogenesis of bladder carcinoma (BCa) using cell line model. The cell lines with low expression of ELF3 showed increased expression of mesenchymal markers and decreased expression of epithelial markers. Immunofluorescence and immunohistochemical analysis of ELF3 showed selective expression in low-grade BCa cell lines and tumor tissues, respectively. We demonstrated that overexpression of *ELF3* in UMUC3, a mesenchymal BCa cell line resulted in reduced invasion and decreased expression of mesenchymal markers. Furthermore, using publicly available data, we found that low expression of *ELF3* was associated with increased risk and poor overall survival rate in BCa. In conclusion, ELF3-modulated reversal of EMT might be a useful strategy in the treatment of bladder cancer.

## Introduction

Bladder carcinoma is incredibly heterogeneous, comprises multiple disease entities with distinct molecular features and biological characteristics.[1] Low-grade non-muscle-invasive (NMI) tumors have high recurrence frequency of 70% and 20% of them progress to muscle invasive tumors (MI).[2] Muscle-invasive tumors are generally diagnosed *de novo* and are involved in metastasis. MI and NMI are two distinct tumors with different clinico-pathological characteristics. NMI carcinoma develop via epithelial hyperplasia whereas MI develop via flat dysplasia and carcinoma *in situ* (CIS).[1] The risk of progression of NMI to MI after 5 years ranges from 6% to 45% [3] with less than 15% of 5 year survival rate.[4] Treatments have not advance beyond cisplatin or cisplatin containing combination chemotherapies for decades.[5] A detailed understanding of bladder carcinoma pathogenesis is an unmet need.

Various studies suggest multifocal occurrence is the characteristic feature of bladder carcinoma. At molecular levels, epithelial to mesenchymal transition (EMT) could play a major role in progression of NMI to MI.[6-9] This transition in cell phenotype is also associated with aggressiveness, recurrence, and metastases of tumors.[10] However, our understanding about the disease pathogenesis is still not evolved comprehensively. EMT is known to be governed by several transcription factors, such as SNAIL, SLUG, ZEB1, ZEB2, and TWIST among others, that transcriptionally repress epithelial markers.[11] These molecules are often referred as EMT transcription factors (EMT-TFs).

Among families of transcription factors, there are increasing evidence for E-twenty six (ETS) transcription factor family members to regulate EMT in cancer.[12-15] ETS family members usually expressed by epithelial cells and are largely implicated in matrix metalloproteinase (MMP) regulation.[16] ETS-related transcription factors are also involved in cell proliferation, adhesion, motility/migration, cell survival, invasion, extravasation, micro-metastasis, establishment and maintenance of distant site metastasis and angiogenesis.[15-17] Member of ETS family, ELF5 (E74 Like ETS Transcription Factor 5) is shown to directly suppress EMT by repressing the transcription of SLUG.[12] ETV1, another member of ETS family also acts to promotes snail expression to induce EMT in gastric cancer.[18] Likewise, ELF3 (E74 like ETS transcription factor 3), also a member of ETS family is restricted to epithelial tissue expression.[19, 20] Higher expression of ELF3 was reported in epithelial rich tissues such as colon, small intestine and negligible expression in epithelial poor tissues such as brain, heart, spleen, thymus and skeletal muscles.[19, 20] Furthermore, expression of ELF3 is negligible in non-epithelial origin cells like lymphocytes, monocytes, and endothelial cells.[20]

Mutation profile studies have identified recurrent anomalies in multiple genes and pathways that are potential key drivers of bladder carcinoma.[21-23] A large-scale genomics study from The Cancer Genome Atlas (TCGA) identified 58 genes as significantly mutated across 412 muscle invasive bladder carcinoma.[24] *ELF3* was reported with frequency of 12% non-synonymous mutation in these tumors. However, the prognostic and functional significance of ELF3 are yet to be explored thoroughly in bladder carcinoma. Here we demonstrate the inverse correlation of ELF3 expression with EMT dynamics along with its association with overall survival. We also studied the effect of *ELF3* overexpression on bladder carcinoma invasiveness using cell based invasion assay.

## Methods

### Cell culture

Five bladder carcinoma cell lines UMUC3, J82, SW780, RT112, T24 were acquired from ATCC and cultured in Dulbecco’s modified eagle medium supplemented with 10% fetal bovine serum, 1% sodium pyruvate and 1% penicillin-streptomycin mixture. Cell lines were grown and maintained in humidified 5% CO_2_ atmosphere at 37°C.

### Transient Transfection

Routinely maintained UMUC3 cells were transfected by piRES-puro-ELF3 (Addgene cat.no. 25728) using X-tremeGENE HP DNA Transfection reagent, Roche, as per manufacture instructions. For western blotting, cells were harvested 72 hours after transfection. For Invasion assay, cells were seeded 24 hours post transfection into boyden chamber.

### Cell invasion assay

The invasive property of UMUC3 was measured using the BD BioCoat™ Matrigel™ Invasion guidelines. Briefly, boyden chamber inserts (Thermo Fisher Scientific, Waltham, MA, USA) were coated with 20 μl 1:10 matrigel and allowed to solidify at room temperature for 1 hour. Cells were seeded in duplicates at 20×10^4^ in 0.5ml serum free media, while the lower chamber contained 10% FBS. Cells were allowed to invade through the porous membrane coated with matrigel at 37°C for 48 hours. Inserts were fixed, stained and photographed in 5 fields per insert. Cell count was performed to determine relative invasion.

### Western blotting

Whole cell extracts from 70-80% confluent cells were prepared using modified RIPA lysis buffer (Merck Millipore, Billerica, MA,) containing protease inhibitors (Roche, Indianapolis, IN,) and phosphatase inhibitors (Thermo Scientific, Bremen, Germany). Western blot analysis was performed on 10% SDS-PAGE using 30 µg protein lysates. After separation proteins were transferred onto nitrocellulose membrane (BioRad) and were hybridized with primary antibodies. Pre blocking in 5% milk for 30 mins was carried out for polyclonal antibodies. Protein bands were visualized using Luminol reagent (Santa Cruz Biotechnology, Dallas, TX,) as per the manufacturer’s instructions. Anti ELF3 antibody was obtained from R&D systems. E-cadherin and slug, antibodies were obtained from Cell Signaling Technology (Cell Signaling Technology, Beverly, MA). Anti N-cadherin and GAPDH were obtained from Abcam. Anti Claudin7 was obtained from Acris antibodies.

### Immunohistochemistry

Bladder carcinoma formalin fixed paraffin embedded tissue sections were obtained following institutional review board approval and informed consent from Kidwai institute of molecular oncology (KMIO/MEC/011/24.November.2016). The sections were deparaffinised and antigen retrieval was carried out using heat-induced epitope retrieval by incubating them for 20 minutes in antigen retrieval buffer (0.01 M Trisodium citrate buffer, pH 6). The quenching of endogenous peroxidases was done by using a blocking solution followed by washes with wash buffer (PBS with 0.05% Tween-20). The sections were incubated with primary antibody overnight at 4°C in a humidified chamber. Anti-ELF3 antibody (R&D systems) was used at 1:200 dilutions. After incubation with the primary antibody, the sections were washed thrice with wash buffer. The slides were incubated with appropriate horseradish peroxidase conjugated goat secondary antibody for 30 minutes at room temperature. Excess secondary antibody was removed using wash buffer followed by addition of DAB substrate. The signal was developed using DAB chromogen (DAKO, Glostrup, Denmark). The immunohistochemical labeling was assessed by an experienced pathologist.

### Immunofluorescent staining

Approximately 3X10^3^ cells were seeded in chamber vials and grown for 48 hours in humidified 5% CO_2_. Cells were fixed with 4% Paraformaldehyde (PFA) (Merk, India) and then permeabilised with 0.5% Triton-X100 (HiMedia, India). Cells were then blocked with 0.1% Triton-X100 in 3% BSA (HiMedia, India). Samples were stained with primary antibody ELF3 (R&D systems), and secondary antibodies - Alexa Fluor^®^ 568-labeled donkey anti-goat IgG (A11057, Invitrogen, CA, USA) and DAPI (D1306, Invitrogen, CA, USA). Images were acquired under inverted epifluorescent microscope (Olympus IX73, USA).

### Computational analysis

Normalized intensities of twenty-six ETS family members were compiled from Earl *et al.,* 2015 for sixteen cell lines.[25] These sixteen cell lines were stratified into group of epithelial cell lines (Epi) (n=9), epithelial or mesenchymal (E/M) (n=2) and mesenchymal (Mes) (n=5) cell lines, as described by Tan *et al.,* 2014.[26] The gene expression matrix was processed using Morpheus (https://software.broadinstitute.org/morpheus/) to generate heatmap with one-minus Pearson correlation clustering approach. SurvExpress database (http://bioinformatica.mty.itesm.mx:8080/Biomatec/SurvivaX.jsp) was used to obtain risk group information for bladder carcinoma.[27] All the 5 different bladder carcinoma datasets were considered for analysis. Risk groups with *p*-values ≤ 0.05 was considered significant.

We also analyzed expression of *ELF3* using web-based platforms such as UALCAN[28] for Kaplan-Meier overall survival analysis to corroborate our cell-line based findings with patient data-sets from The Cancer Genome Atlas (TCGA).

### Computation of EMT score

Meta-cohort of bladder carcinoma dataset Affymetrix U133A platform was compiled in Tan *et al*., 2014.[26] Briefly, bladder carcinoma from GSE31684, GSE7476, and GSE5287 were downloaded from GEO. The data were RMA-normalized and combined using ComBat.[29] Clinical and processed RNA-seq data of TCGA bladder carcinoma cohort was downloaded from GDAC (version 2016_01_28) (Broad2016). The FPKM values of *ELF3* were extracted for downstream analysis.

## Results

### Expression pattern of ELF3 and ETS family members

EMT score computed by Tan *et al.,* 2014 classified RT4, UMUC9, HT1197, HT1376, BC16.1, CUBIII, SCABER, UMUC6, PSI, 5637, KU7, as epithelial cell lines.[26] HU456 and KK47 were stratified as cell lines showing both epithelial and mesenchymal characteristics.[26] Whereas 253JBV, T24, TCCSUP, J82, UMUC13 and UMUC3 showed most mesenchymal characteristics. We further utilized same classification to assess expression pattern of ETS family members from microarray data provided by Earl *et al.,* 2015 across 40 bladder carcinoma cell lines (Supplementary table S1).[25] We observed that *ELF2, ELF1, ETS2, ERF, ELF4, ETV4, ETS1, SPDEF, ELK1, FEV, ETV1* were highly expressed genes across cell lines. Whereas *FL1, ELK4, ETV3, ERG* and *ETV2* showed relatively low expression across cell lines (Figure 1A). We observed *ELF3* expression was relatively low in cell lines characterized as mesenchymal (T24, TCCSUP, J82, UMUC13 and UMUC3) and high in cell lines characterized as epithelial (RT4, UMUC9, HT1197, HT1376, PSI and 5637) (Figure 1B). Other epithelial cell lines SCABER and UMUC6 presented moderate expression of *ELF3* whereas UMUC14 exhibited low expression of *ELF3*. *ELF3* expression difference was assessed using non-parametric unpaired Mann-Whitney test and we observe that it was significantly higher in epithelial cell lines compared to mesenchymal cell lines with *p*-value of 0. 0280.

**Figure 1.**
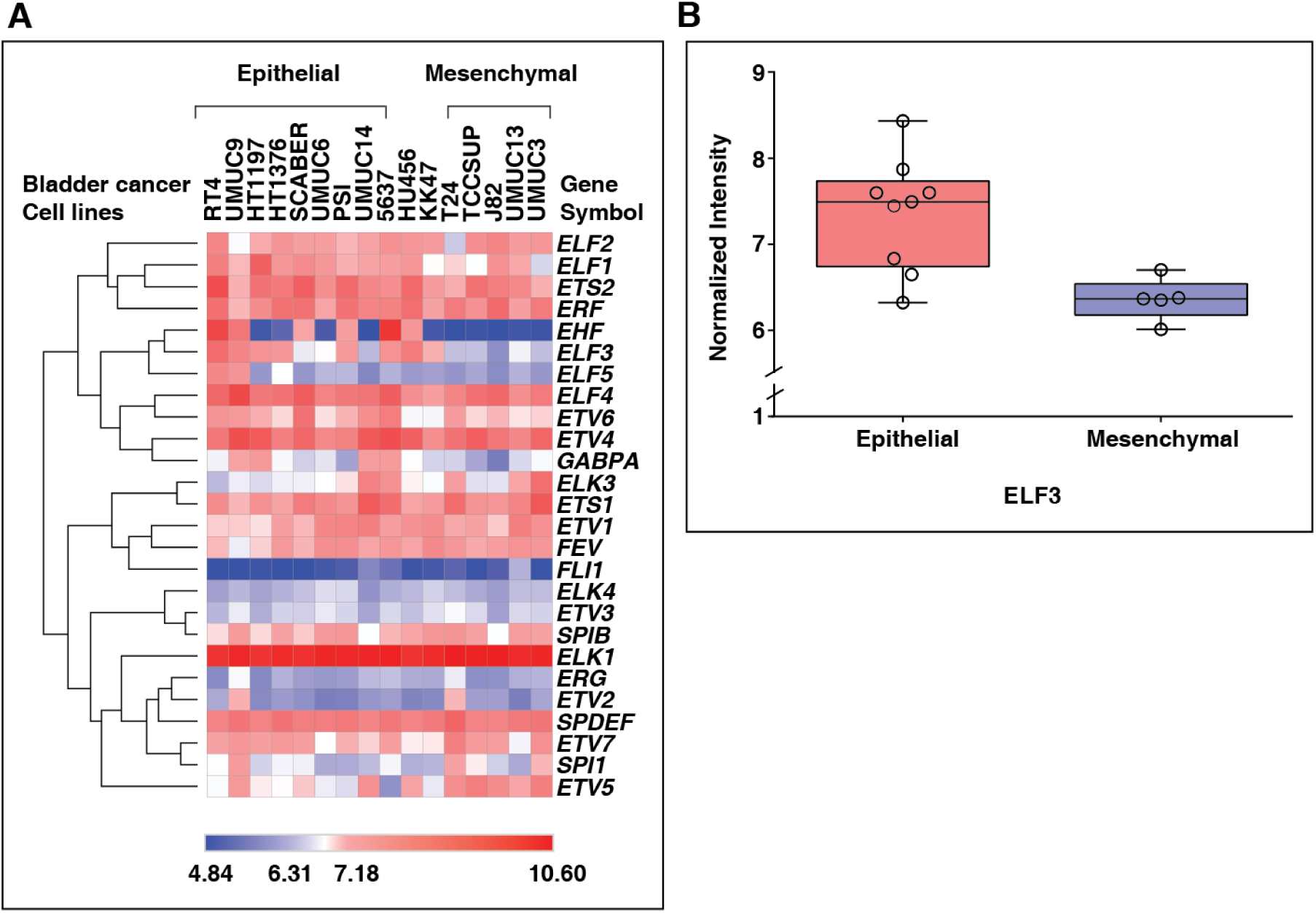
Expression profile of ETS family in bladder carcinoma cell lines. **(A)** Heatmap representing normalized intensities of ETS family from epithelial (RT4, UMUC9, HT1197, HT1376, BC16.1, CUBIII, SCABER, UMUC6, PSI, 5637, KU7), mixed (HU456 and KK47) and mesenchymal (253JBV, T24, TCCSUP, J82, UMUC13, UMUC3) tumor derived cell lines reported using microarray analysis **(B)** Relatively low expression of ELF3 in mesenchymal compared to epithelial tumor derived cell lines

### ELF3 is down regulated in cell lines derived from high grade tumors

To confirm the expression pattern of ELF3, western blot was performed across 5 bladder carcinoma cell lines. These cell lines were established from low-grade tumor to high-grade tumors (Supplementary table S2). As shown in figure 2A-E and figure 3A, cell lines derived from low grade tumor, SW780 and RT112 expressed ELF3 while other cell lines T24 and J82 derived from high grade tumors and also characterized as most mesenchymal cell lines along with UMUC3 showed negligible expression of ELF3. We also observed varied localization pattern of ELF3 with respect to grade of the cell lines. ELF3 was localized in nucleus and cytoplasm in SW780 and RT112 whereas in T24 and J82, ELF3 was localized in negligible amounts in cytoplasm and absent in nucleus.

**Figure 2.**
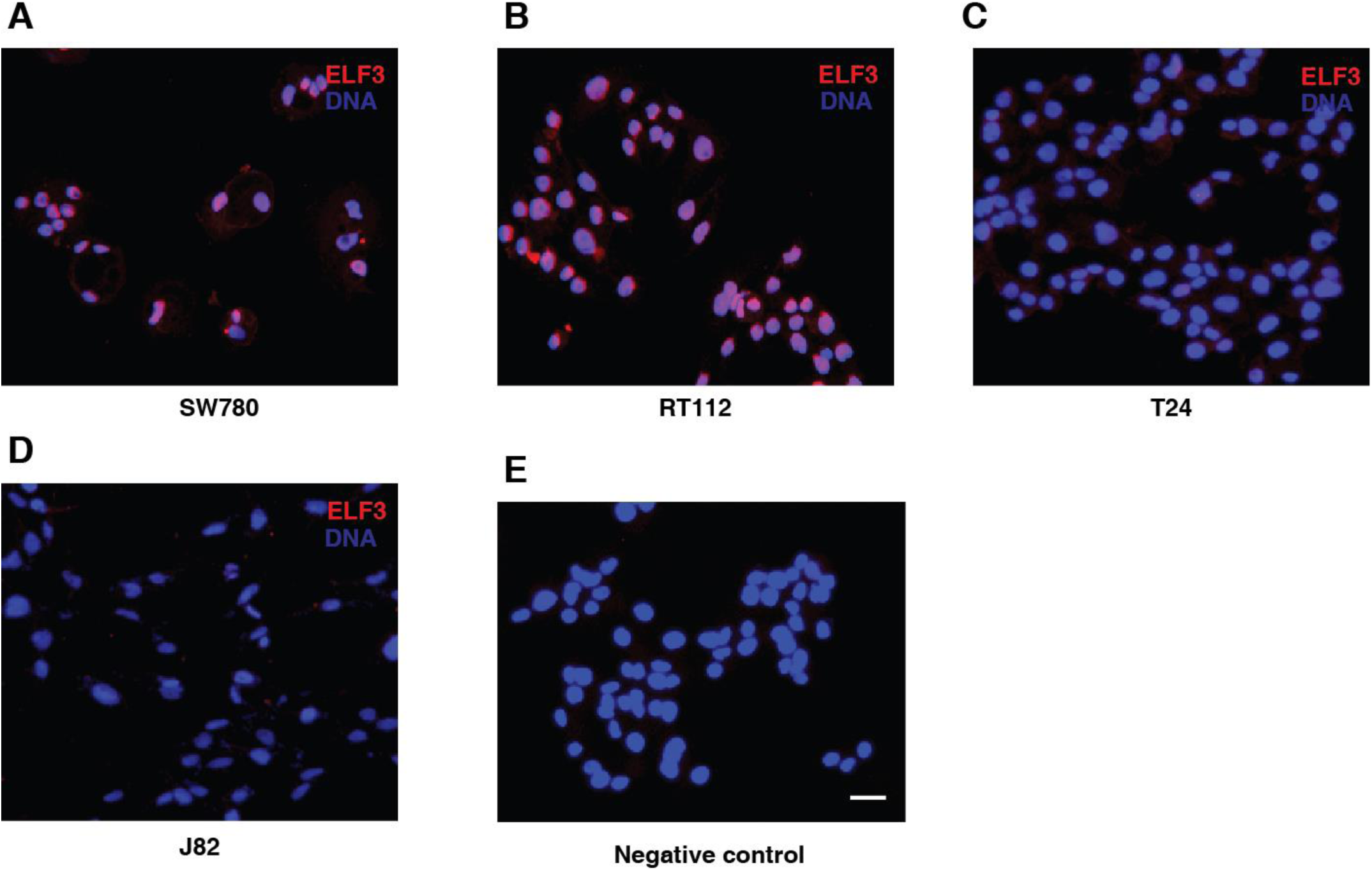
Subcellular localization of ELF3 in bladder carcinoma cell lines. **Immunofluorescence staining showed cytoplasmic and nuclear localization of ELF3 in (A)** SW780 **(B)** RT112; and cytoplasmic in **(C)** T24, **(D)** J82. **(E)** Negative control

**Figure 3.**
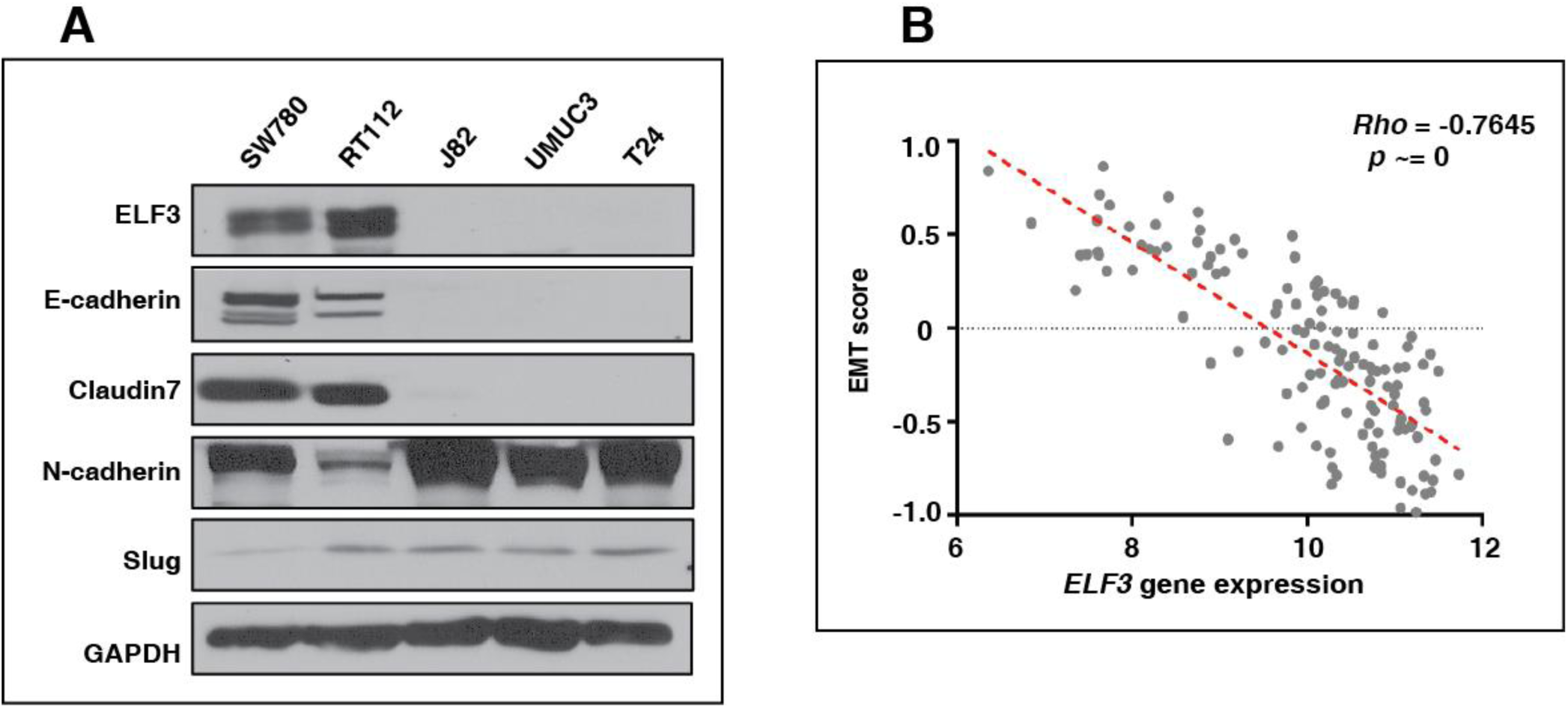
Association of ELF3 expression with EMT signature. **(A)** Western blot analysis depicting higher expression of epithelial markers E-cadherin and Claudin7 in SW780 and RT112; and higher expression of mesenchymal markers N-cadherin and slug in J82, UMUC3 and T24 **(B)** Spearman’s inverse correlation of EMT signature with ELF3 expression in bladder carcinoma patient data procured from Tan *et al.*

### ELF3 expression is associated with epithelial to mesenchymal transition in bladder carcinoma cells

To study the functional role of ELF3 in EMT process, western blot analysis was performed in bladder carcinoma cell lines against the expression pattern of various EMT markers. We observed low expression of ELF3 correlated with low expression of epithelial markers E-cadherin and Claudin7 in high grade bladder carcinoma cell lines, J82, T24 and mesenchymal cell line UMUC3. Whereas expression of mesenchymal markers such as N-cadherin and SLUG were higher in high grade compared to low grade bladder carcinoma cell lines (Figure 3A). Expression of ELF3 might be involved in negative regulation of EMT in bladder carcinoma. To further validate our findings, we compared expression of ELF3 with the generic EMT signature derived from bladder carcinoma patient data sets as described by Tan *et al.*, 2014.[26] This scoring method was developed to quantitatively estimate the EMT phenotype across clinical samples as well as cell lines using transcriptomics. Based on this *in-silico* analysis, we identified that low *ELF3* expression significantly correlated (Rho = -0.7645) with higher mesenchymal phenotype in bladder carcinoma patients (Figure 3B).

### Over expression of *ELF3* decreases invasion in mesenchymal bladder carcinoma cell line UMUC3

Expression of ELF3 was downregulated across high-grade bladder carcinoma cell lines. We further sought to identify whether *ELF3* overexpression can potentially revert the invasiveness in UMUC3 cell line. We observed significant decreased invasiveness of UMUC3 (Figure 4A). *ELF3* over expression also showed marginal decrease in expression of mesenchymal markers like N-cadherin and SLUG, however there was no change in the expression of epithelial markers (Figure 4B) as confirmed by western blot.

**Figure 4.**
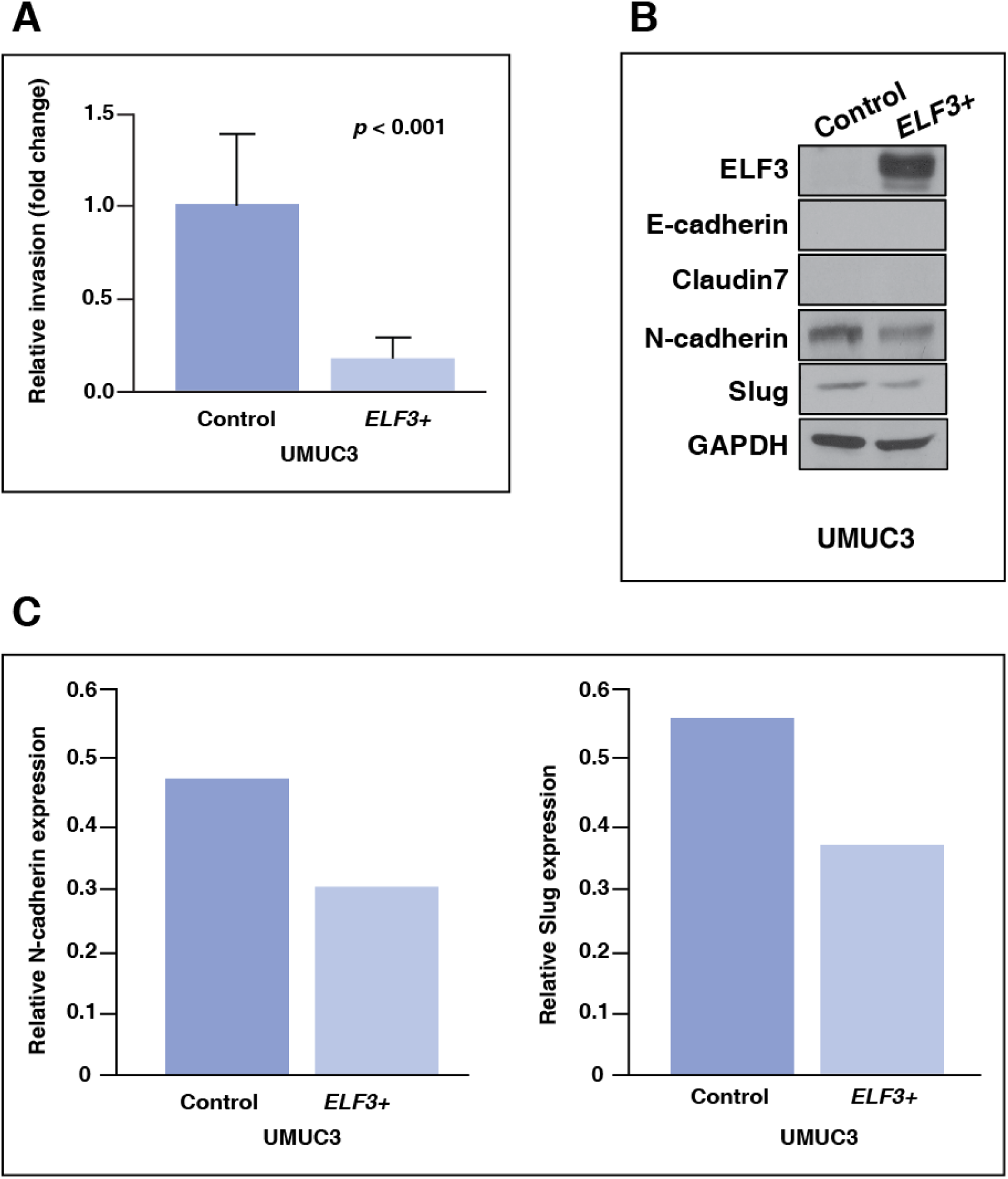
Overexpression of ELF3 reduces cell invasion in-vitro. **(A)** Transient transfection of ELF3 in UMUC3 lead to decreased invasiveness **(B, C)** Western blot and imageJ analysis indicates lower expression of mesenchymal markers and unchanged expression of epithelial markers.

### Immunohistochemistry revealed higher expression of ELF3 in low grade compared to high grade tumors

To further investigate the expression pattern of ELF3, we performed immunohistochemical staining on bladder tumors. Patient presented with low grade carcinoma showed unchanged ELF3 expression in tumor compared to adjacent normal (Figure 5A, B). Whereas patient presented with high grade carcinoma showed decreased level of ELF3 expression in tumor compared to adjacent normal (Figure 5C, D). We observed considerable difference in the ELF3 expression between low grade and high grade tumors. In addition, we also observed decreased ELF3 expression in mesenchymal cells compared to the epithelial cells in the high grade tumor (Figure 5E). Our result suggests that ELF3 expression is higher in low grade compared to high grade tumors.

**Figure 5.**
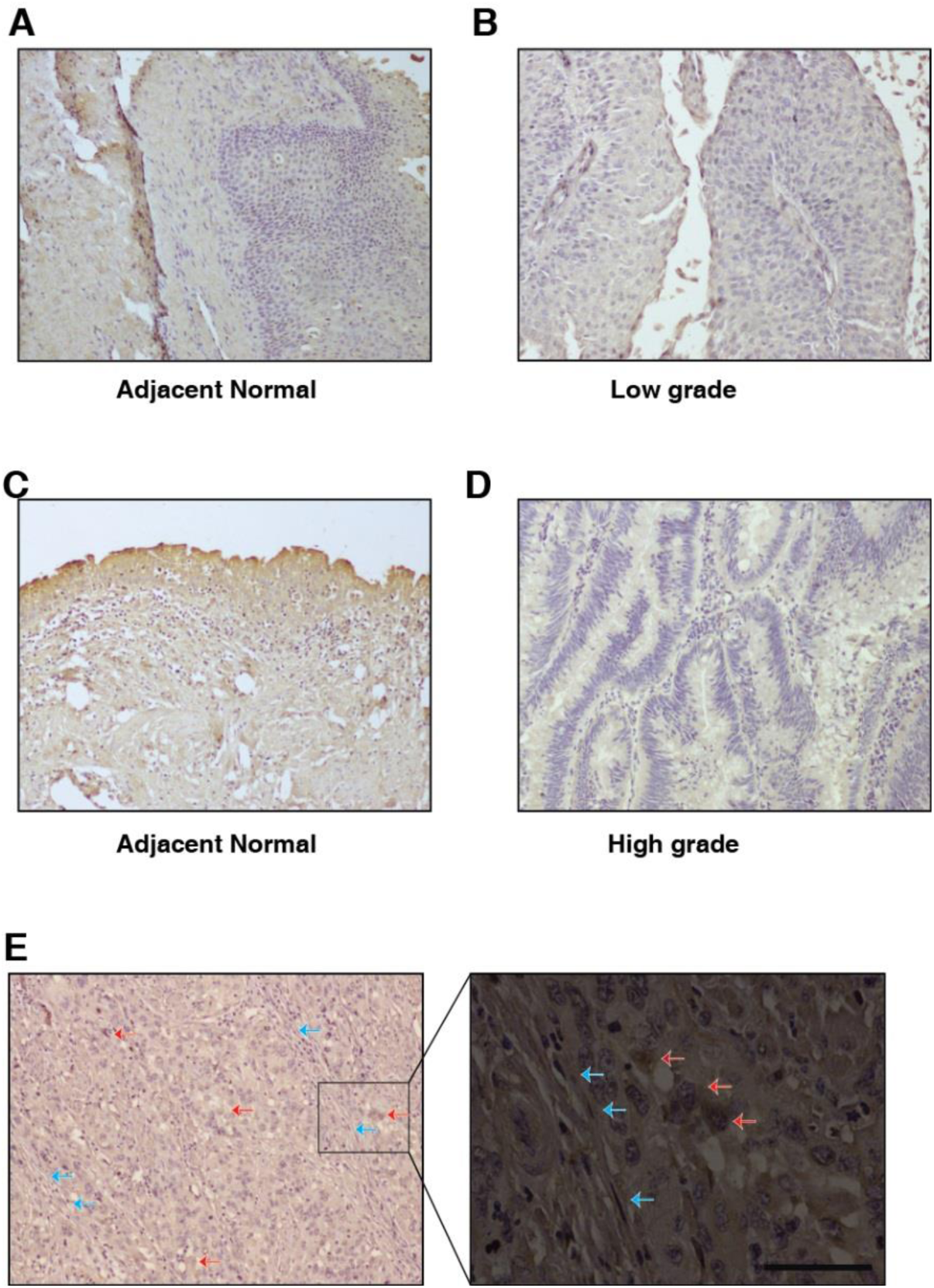
ELF3 expression in different grades of urothelial tumors. Immunohistochemical staining show comparable expression of ELF3 in **(A)** adjacent normal and **(B)** tumor of low grade urothelial carcinoma; whereas decreased expression from **(C)** adjacent normal to **(D)** tumor of high grade urothelial carcinoma. **(E)** IHC image depicts decreased expression of ELF3 in mesenchymal cells compared to epithelial cells (Blue arrows: mesenchymal cells. Red arrows: epithelial cells)

### Association of ELF3 downregulation with high risk and overall poor survival in bladder carcinoma

To assess potential risk associated with *ELF3* expression in bladder carcinoma we have used five published studies hosted on an online biomarker validation tool SurvExpress.[27] These datasets were classified into low risk and high risk groups, and *p*-value was calculated using student *t* test. From the aforementioned five data-sets, four i.e. bladder urothelial carcinoma TCGA (n=54), GSE31684 (n=93), GSE5287 (n=30) and BLCA-TCGA-Bladder urothelial carcinoma (n=390) presented that low gene expression of *ELF3* was significantly correlated to high risk with hazard ratios of 1.28, 1.08, 0.87, 1.5 respectively (Supplementary figure 1)

To investigate the association between *ELF3* expression and overall survival, we analysed publicly available datasets using Kaplan-Meier method using UALCAN.[28] Survival analysis revealed patients with low *ELF3* expression (n=304) had poor overall survival compared to patients with high expression (n=102, log rank *p* value = 0.12) (Supplementary figure 2). Results suggest that low expression of *ELF3* is correlated with adverse outcomes.

## Discussion

ELF3 also named as ESX, ERT and ESE-1 is an epithelial-restricted member of the ETS transcription factor family, and involved in wide range of cellular processes such as differentiation, and inflammatory response.[28, 30] It is one of the critical regulators involved in urothelial cyto-differentiation in normal human urothelial (NHU) cell line.[30] Expression of ELF3 is reported in nucleus as well as in cytoplasm.[31] ELF3 positively regulates expression of TGFβR-II[32] and is critically important in regulation of intestinal epithelial differentiation in mice.[33] Mounting evidence suggests that transcription factors play a key role in cancer progression.[34] Like most transcription factors, ETS proteins act as both oncogene and tumor suppressors.[35] It has been also reported that ELF3 controls the transcription of various genes that are involved in extracellular matrix remodeling like collagenase-1, matrix metalloproteinases MMP1 and MMP13, further enhancing the process of cartilage remodeling or degradation and tumorigenesis. [36, 37]

We interrogated *ELF3* expression in TCGA datasets using web-based resource UALCAN to corroborate our findings from cell line in clinical samples. UALCAN is implemented in PERL-CGI to enable web-based gene expression analysis in various stages and molecular subtypes of 31 different cancers.[28] We observed there is lower expression of ELF3 in basal/squamous (n=142) bladder carcinoma subtype compared to luminal (n=246) and normal tissues (Supplementary figure 3). Squamous differentiated urothelial carcinoma are associated with advanced stages of carcinoma[38] and basal like urothelial carcinoma are characterized as non-papillary aggressive tumors with higher rates of lymph node and distant metastases compared to non-basal like urothelial carcinoma.[39]

To further understand the role of ELF3 in EMT, we investigated the expression of ELF3 in bladder carcinoma cell lines. We observed the level of ELF3 was specifically low in mesenchymal cells. Immunohistochemical analysis in low grade and high grade bladder tumors suggests decrease in the expression of ELF3 from low grade to high grade. In addition, the expression of epithelial markers were observed low in high grade cell lines and inversely correlated with the expression of mesenchymal markers. It suggests that ELF3 might be involved in regulation of EMT. In this study, we identified similar expression pattern of ELF3 and Claudin7 as reported by Kohno *et al.*, which suggests ELF3 is indeed involved in the formation of epithelial structure in bladder cancer.[40]

Various studies have indicated localization of ELF3 affects its function. ELF3 mediates the translocation of snail and β-catenin to cytoplasm to induce EMT.[31],[41] ELF3 is also shown to induce anchorage-independent growth and transformation of MCF-12A cell line via cytoplasmic mechanism.[42] In our study, we observed that ELF3 was specifically localized in cytoplasm in high grade bladder carcinoma cell lines T24 and J82 which might be a mechanism that imparts mesenchymal nature to these cell lines. We also observed the cytoplasmic expression of ELF3 in low-grade tumor sections. However, the exact reason for difference in localization in bladder carcinoma cell lines is not known yet.

We further sought to understand the process of bladder carcinoma progression. We checked the gain of function of *ELF3* in bladder cell lines where negligible expression of ELF3 was observed. We found that upon *ELF3* transfection in mesenchymal cell line UMUC3, the invasive ability of the cell line was decreased significantly. Transient transfection also showed a slight decrease in expression of mesenchymal markers N-cadherin and Slug. Subsequently we also checked the effect of *ELF3* overexpression and EMT modulation in J82 and T24; however, these cell lines were very difficult to transfect and we could not get more than 10% transfection efficiency.

We observed that low expression of *ELF3* correlated significantly with poor survival in bladder carcinoma patients. Our meta-analysis also suggested that patients are at higher risk with low *ELF3* expression in 4 out of 5 bladder carcinoma published data-sets as per SurvExpress.[27] Low expression of *ELF3* is also reported to have effect on patient’s overall survival and *ELF3* over expression could also be used to develop novel therapeutic regimens and could be implicated to improve bladder carcinoma patient survival. Our study delineates the molecular mechanism of ELF3 mediated EMT in aggressive bladder tumors.

## Conclusions

Epithelial to mesenchymal transition phenomenon majorly contributes to metastasis and cancer progression, however the molecular mechanisms underlying this transition remains to be fully understood. Our study revealed the role of ELF3 in the cellular plasticity of bladder carcinoma. This study for the first time demonstrates that expression of ELF3 negatively correlates with expression of epithelial to mesenchymal transition markers and ELF3-modulated reversal of EMT can be a valuable strategy in treatment of bladder carcinoma.

## Supporting information

## Acknowledgements

PK is a recipient of the Ramanujan Fellowship awarded by Department of Science and Technology (DST), Government of India. KG is a recipient of Senior Research Fellowship from University Grants Commission (UGC), Government of India. KP is a recipient of Senior Research Fellowship from Council of Scientific Industrial Research, Government of India.

## Supplementary Figure Legends

**Supplementary Figure 1. Association of ELF3 expression with risk groups in bladder carcinoma** Downregulation of ELF3 is observed to have significant association with high risk in datasets (A) Bladder urothelial carcinoma TCGA (B) GSE31684 (C) GSE5287 (D) BLCA-TCGA, 2016 and low risk in (E) GSE13507

**Supplementary Figure 2. Low expression of ELF3 is associated with overall poor survival in bladder carcinoma patients**

**Supplementary Figure 3. Relative low expression of ELF3 in basal squamous subtype compared to other subtypes of bladder carcinoma**

**Supplementary Table Legends**

**Supplementary Table S1: Gene expression of ETS family members across bladder carcinoma cell lines from Earl et al., 2015**

**Supplementary Table S2: Bladder carcinoma cell lines and their origin**

