## Supplementary material for "E74 like ETS transcription factor 3 (ELF3) is a negative regulator of epithelial-mesenchymal transition in bladder carcinoma"

### Supplementary Figure 1

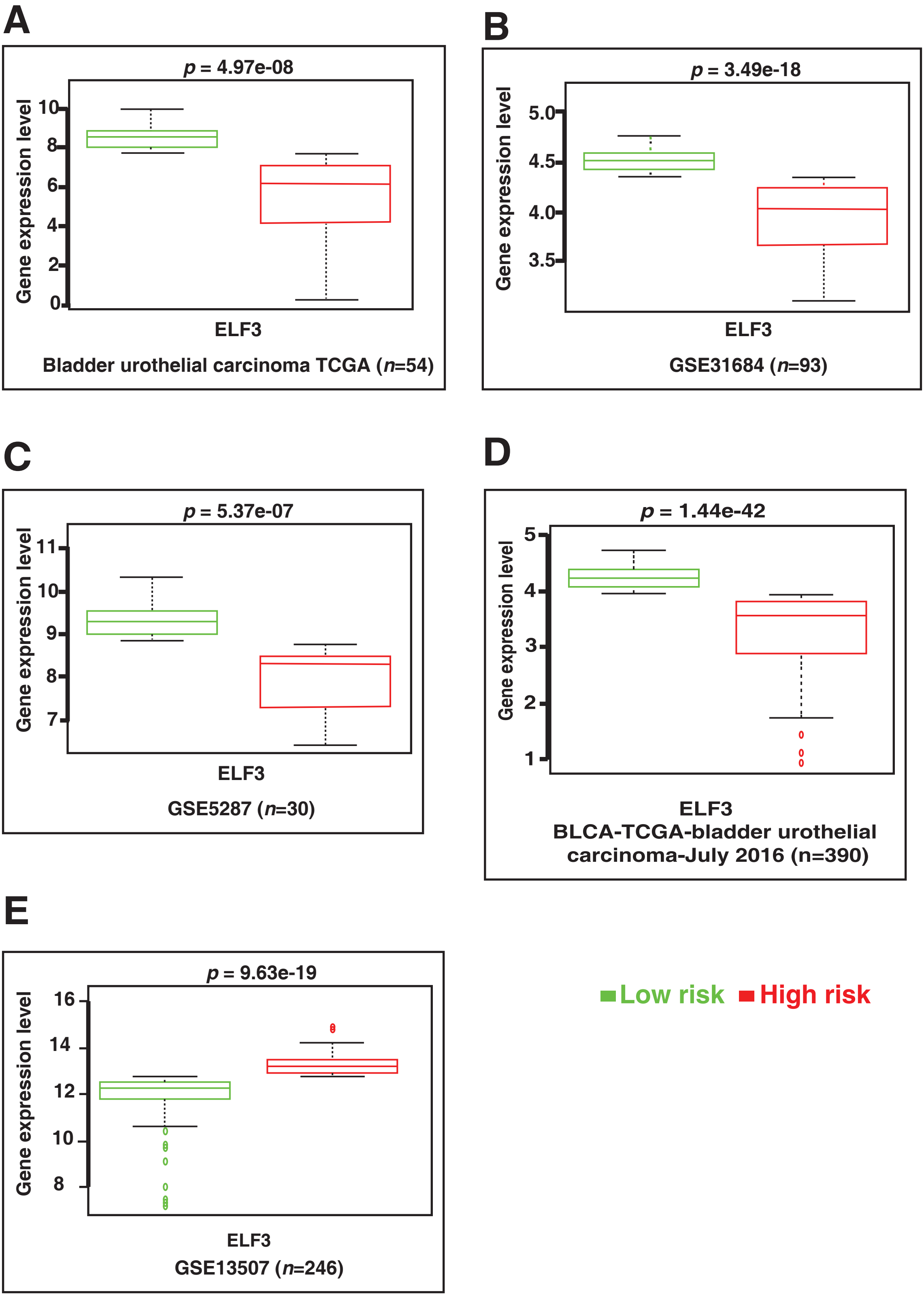

### Supplementary Figure 2

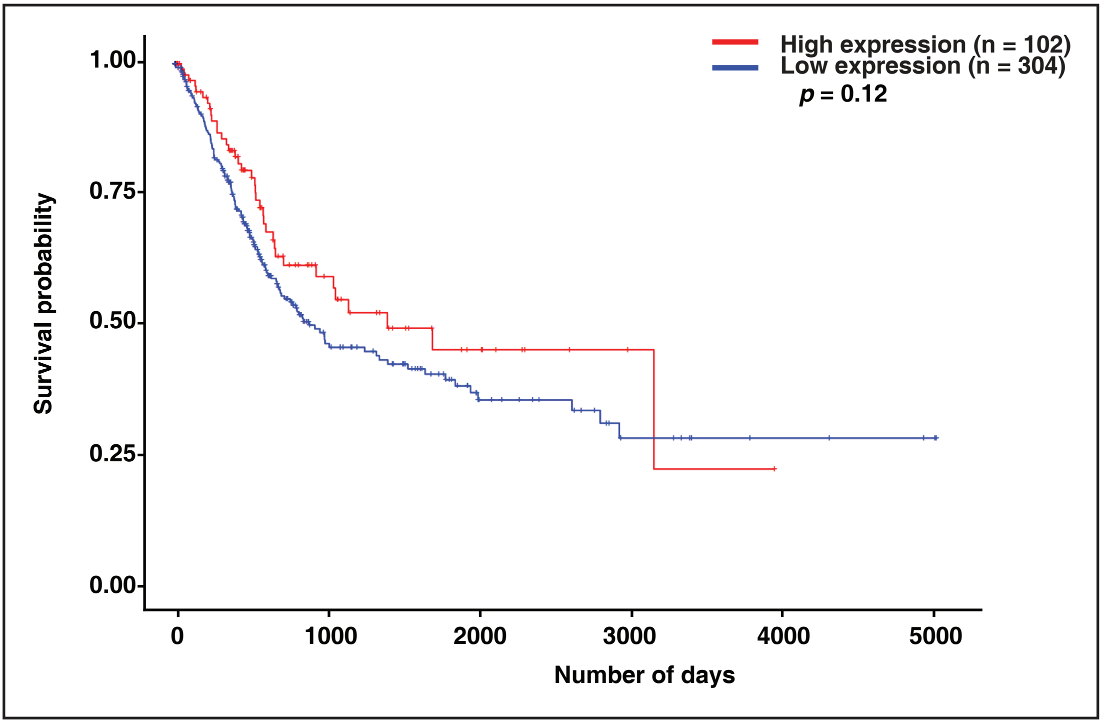


### Supplementary Figure 3

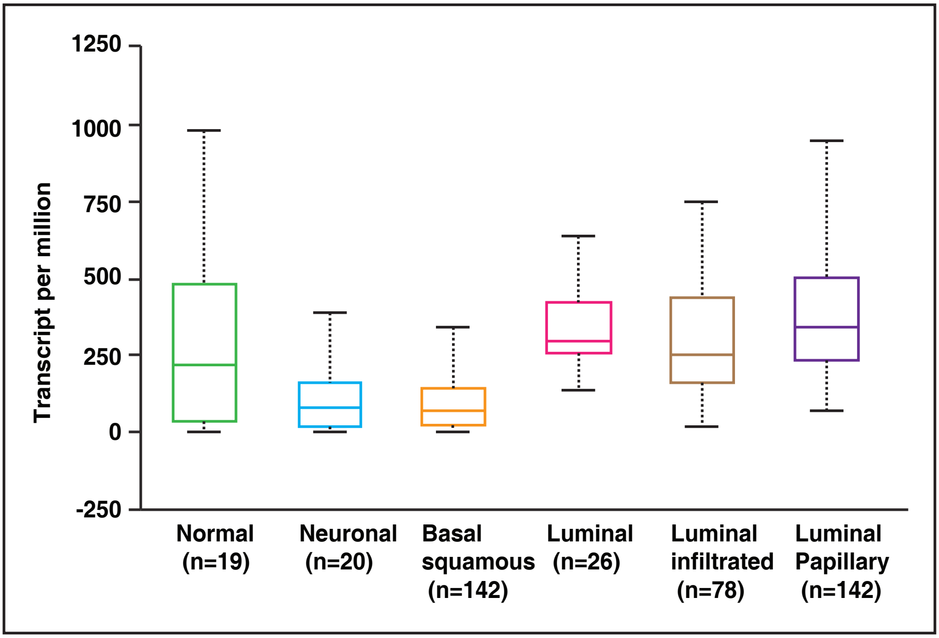

**Gondkar *et al.* , 2018. E74 like ETS transcription factor 3 (ELF3) is a negative regulator of epithelial-mesenchymal transition in bladder carcinoma.**

**Supplementary Table 1: Expression values of ETS family members across bladder cancer cell lines from Earl et al., 2015**

| | Epithelial | | | | | | | | | E/M | | Mesenchymal | | | | |
|---|---|---|---|---|---|---|---|---|---|---|---|---|---|---|---|---|
| **Gene Symbol** | **RT4** | **UM-UC-9** | **HT1197** | **HT1376** | **SCABER** | **UM-UC-6** | **PSI** | **UM-UC-14** | **5637** | **HU456** | **KK47** | **T24** | **TCCSUP** | **J82** | **UM-UC-13** | **UM-UC-3** |
| *EHF* | 9.280510411 | 8.165978396 | 5.17145688 | 5.63199306 | 7.243589172 | 5.325919342 | 7.352115149 | 4.918334472 | 9.664578217 | 7.529522343 | 5.217824181 | 4.846882534 | 5.041505841 | 4.883326446 | 5.122860301 | 5.042836168 |
| *ELF1* | 8.00331677 | 7.100471758 | 8.638688448 | 7.484786645 | 7.741196006 | 7.515717354 | 7.142908859 | 7.261166313 | 7.347632587 | 7.480473475 | 6.838619888 | 7.002538083 | 6.803378021 | 7.640834356 | 7.187296412 | 6.525301927 |
| *ELF2* | 7.834959177 | 6.788871273 | 7.16716659 | 7.570105831 | 7.316847866 | 7.345660087 | 7.120925328 | 7.26275268 | 7.640824429 | 7.482546175 | 7.405103775 | 6.426237965 | 7.69219112 | 7.873738228 | 7.383916721 | 7.479071214 |
| *ELF3* | 8.432504096 | 7.870296868 | 7.598686293 | 7.493140136 | 6.6491018 | 6.833186624 | 7.449845149 | 6.323144853 | 7.597853264 | 8.075227198 | 7.144713707 | 6.366725902 | 6.37775602 | 6.010953171 | 6.701627681 | 6.35176381 |
| *ELF4* | 8.499626746 | 9.19363403 | 8.166980849 | 8.268925826 | 8.755128644 | 7.910467008 | 8.143287853 | 8.203251966 | 8.679796502 | 7.682598847 | 7.341361335 | 7.883046645 | 8.316786285 | 8.494937749 | 7.718962532 | 8.056945122 |
| *ELF5* | 7.853993234 | 7.572222198 | 6.078863211 | 6.80157422 | 6.066922413 | 6.301878245 | 6.314321443 | 5.95536808 | 6.260726828 | 6.099315334 | 6.131457701 | 6.055468815 | 6.177016224 | 5.988157817 | 6.27143997 | 6.073845404 |
| *ELK1* | 9.720478711 | 10.05133 | 9.695362901 | 10.02123326 | 9.809500684 | 10.17618205 | 10.05866698 | 10.07508901 | 10.30353645 | 9.741972111 | 10.020742 | 10.60367897 | 10.23031194 | 10.47415067 | 9.991390346 | 10.16928095 |
| *ELK3* | 6.326153745 | 6.681428506 | 6.540380258 | 6.691207992 | 6.696625806 | 6.820075176 | 6.940562243 | 7.976984597 | 7.649794955 | 6.879290087 | 6.772692473 | 7.36665083 | 6.588286705 | 6.612670026 | 7.215880421 | 8.346807509 |
| *ELK4* | 6.134467657 | 6.296707572 | 6.154243447 | 6.277696883 | 6.374829103 | 6.554855818 | 6.531211073 | 6.044942728 | 6.234553402 | 6.318445656 | 6.496274241 | 6.341983085 | 6.197007731 | 6.107559489 | 6.344964432 | 6.34227704 |
| *ERF* | 8.295221926 | 7.117536767 | 7.755639443 | 8.285633633 | 8.286444707 | 7.238007115 | 8.268372373 | 7.896649539 | 7.951643894 | 8.350539971 | 7.307025452 | 8.054229934 | 7.647642031 | 8.452655847 | 7.344394527 | 8.054753776 |
| *ERG* | 5.971979756 | 6.758245646 | 5.9659786 | 6.27621043 | 6.111867837 | 6.081350748 | 6.107384289 | 6.325721597 | 6.362535537 | 6.217071098 | 6.201386774 | 6.663586476 | 6.008454402 | 6.026068352 | 6.24655324 | 6.283933715 |
| *ETS1* | 7.733962572 | 7.112642263 | 7.655086255 | 7.256680344 | 8.047958947 | 7.789260042 | 7.580424677 | 8.80608728 | 8.389674654 | 7.250137833 | 7.408025774 | 8.401014474 | 7.584896413 | 7.521073433 | 7.872551691 | 8.87069066 |
| *ETS2* | 9.082279179 | 7.136382297 | 8.2411605 | 8.114632631 | 8.659961467 | 7.709268326 | 8.446443453 | 7.825253503 | 7.683178324 | 8.455457015 | 7.378552829 | 7.156763804 | 8.287549181 | 8.002664704 | 7.808349905 | 7.147937702 |
| *ETV1* | 7.015319573 | 7.026784794 | 6.94744464 | 7.327426117 | 7.107201191 | 7.837285793 | 7.927924133 | 8.000179335 | 7.372402273 | 7.65441623 | 7.720685847 | 7.176236872 | 7.310102197 | 7.031931466 | 8.016509006 | 7.645134647 |
| *ETV2* | 6.190377112 | 7.138736187 | 5.980086278 | 6.095582385 | 6.028371122 | 5.878394499 | 5.90113757 | 6.065153261 | 6.115031308 | 5.925249257 | 5.967566965 | 7.113337728 | 6.121014742 | 6.124080861 | 5.87935437 | 6.172778077 |
| *ETV3* | 6.295970969 | 6.672012621 | 6.229218847 | 6.491141575 | 6.473725434 | 6.641928314 | 6.505017986 | 6.172162463 | 6.528758302 | 6.360622699 | 6.597202071 | 6.764833199 | 6.540410882 | 6.133635233 | 6.537102094 | 6.490383221 |
| *ETV4* | 8.134139545 | 9.062838807 | 8.826156545 | 8.001896884 | 8.529708564 | 7.818729175 | 7.955199096 | 8.85795782 | 9.067565489 | 8.650107119 | 7.710935837 | 8.261646769 | 8.663652834 | 8.148259584 | 7.850151271 | 8.514561702 |
| *ETV5* | 6.774902564 | 7.458468884 | 6.901655911 | 6.833257519 | 7.045568813 | 6.676973097 | 6.557934774 | 7.63606444 | 6.02543561 | 7.262839162 | 6.641616109 | 7.69556684 | 8.021737955 | 7.721872371 | 7.190017351 | 7.996223931 |
| *ETV6* | 7.584124977 | 7.437315078 | 7.148885531 | 6.982552739 | 8.280586044 | 6.947605195 | 7.13034858 | 7.708891136 | 8.043313189 | 6.743338498 | 6.730322903 | 7.384049963 | 6.984160102 | 7.112334989 | 6.940127445 | 7.006409165 |
| *ETV7* | 7.286897208 | 7.491637919 | 7.316012688 | 7.445003328 | 7.34476858 | 6.819868075 | 7.143485381 | 7.000612339 | 7.36850596 | 6.889931951 | 6.915804588 | 7.995271413 | 7.428679669 | 7.370442015 | 6.720909605 | 7.66770199 |
| *FEV* | 7.103176316 | 6.679170501 | 7.012455251 | 7.458917821 | 7.1306337 | 7.701816508 | 7.695155084 | 7.233006109 | 7.513702158 | 7.23702657 | 7.799316277 | 7.500714421 | 7.169811059 | 7.3358926 | 7.606813037 | 7.488363565 |
| *FLI1* | 4.879552002 | 4.846297439 | 5.04547147 | 4.848663215 | 4.858792947 | 5.184916964 | 5.404227963 | 5.951295449 | 5.767862038 | 5.04074577 | 5.242009639 | 5.572190265 | 5.002513366 | 5.45263896 | 6.255831993 | 4.900730975 |
| *GABPA* | 6.699163 | 7.257457726 | 7.472952817 | 6.742741141 | 6.469871221 | 6.62258402 | 6.149226772 | 7.356453055 | 7.425126783 | 6.797510867 | 6.498516824 | 6.488055062 | 6.178263425 | 5.862531003 | 6.527594245 | 6.765765604 |
| *SPDEF* | 7.941050744 | 8.222295729 | 7.920914699 | 8.234551448 | 8.006085668 | 8.036340387 | 8.117126348 | 7.874181861 | 8.159552537 | 7.928094702 | 8.088754355 | 8.54398369 | 7.92190748 | 7.92417795 | 8.088864604 | 8.19040829 |
| *SPIB* | 6.953626084 | 7.429959244 | 6.978554307 | 7.2972571 | 7.037403232 | 7.440236796 | 7.494440054 | 6.804637615 | 7.118987853 | 7.212038209 | 7.503314543 | 7.590447097 | 7.28881638 | 6.799620315 | 7.353054877 | 7.311690727 |
| *SPI1* | 6.792251626 | 7.409428023 | 6.49520546 | 6.718350199 | 6.696964545 | 6.200107586 | 6.212811974 | 6.338096844 | 6.729846069 | 6.252886873 | 6.291412889 | 7.288474789 | 6.902574531 | 6.473992773 | 6.191455552 | 7.114393224 |

**Gondkar et al., 2018. E74 like ETS transcription factor 3 (ELF3) is a negative regulator of epithelial-mesenchymal transition in bladder carcinoma.**

**Supplementary Table 2: Bladder carcinoma cell lines and their origin**

| Cell line | Origin of the cell line | Grade | Reference |
|---|---|---|---|
| SW780 | Transitional cell carcinoma | Grade 1 | Fogh J., Natl Cancer Inst Monogr., 1978 |
| RT112 | Transitional cell carcinoma | Grade 2 | Marshall CJ., *et al* ., J Natl Cancer Inst. 1977 |
| T24 | Transitional cell carcinoma | Grade 3 | Bubenik J., *et al* .,Int J Cancer., 1973 |
| UMUC3 | Transitional cell carcinoma | Not reported | Grossman HB., *et al* ., J Urol., 1986 |
| J82 | Transitional cell carcinoma | Grade 3 | O'Toole C., *et al* ., Br J Cancer., 1978 |
